## Supplementary Material for "Time-resolved NMR monitoring of tRNA maturation"

**Content:**


| | |
|---|---|
| Supplementary Table S1 to S3 | page 2-4 |
| Supplementary Figures S1 to S11 | pages 5-15 |
| Supplementary References | page 16 |

**Supplementary Table S1: Overview of the 14 modifications present in yeast cytoplasmic $tRNA^{Phe}$.**

| modification | enzyme | gene | partner protein | localization |
|---|---|---|---|---|
| **$m^2G10$** | Trm11 | *trm11* | Trm112 | cytoplasm |
| **D16** | Dus1 | *dus1* | - | nucleus |
| **D17** | Dus1 | *dus1* | - | nucleus |
| **$m^2_2G26$** | Trm1 | *trm1* | - | nucleus |
| **Cm32** | Trm7 | *trm7* | Trm732 | cytoplasm |
| **Gm34** | Trm7 | *trm7* | Trm734 | cytoplasm |
| **yW37** | Trm5, TYW1-4 | *trm5, tyw1-4* | - | nucleus/cytoplasm[a] |
| **Ψ39** | Pus3 | *pus3* | - | nucleus |
| **$m^5C40$** | Trm4 | *trm4* | - | nucleus |
| **$m^7G46$** | Trm8 | *trm8* | Trm82 | nucleus |
| **$m^5C49$** | Trm4 | *trm4* | - | nucleus |
| **$m^5U54$ (T54)** | Trm2 | *trm2* | - | nucleus[b] |
| **Ψ55** | Pus4 | *pus4* | - | nucleus |
| **$m^1A58$** | Trm61 | *trm61* | Trm6 | nucleus |

Modifications are listed in the first column, the catalytic enzyme responsible for their introduction in the second column and the corresponding genes in the third column. Partner proteins, when present are listed in the fourth column. The known sub-cellular localizations (nucleus or cytoplasm) of the enzymes are given in the fifth column.

[a] The complex modification yW37 (wybutosine37) requires five enzymes for its biosynthesis, *i.e.* Trm5 and TYW1-4, which are found in the nucleus for Trm5 and cytoplasm for TYW1-4 (1).

[b] from indirect evidence (2,3).

**Supplementary Table S2: Strains used in this study.**

| strain | genotype | source |
|---|---|---|
| BY4741 | *MATa his3-Δ1 leu2-Δ0 met15-Δ0 ura3-Δ0* (wild type) | Euroscarf |
| BY4741-*dus1Δ* | BY4741, *YML080w::kanMX* | Euroscarf |
| BY4741-*pus4Δ* | BY4741, *YNL292w::kanMX* | Euroscarf |
| BY4741-*trm1Δ* | BY4741, *YDR120c::kanMX* | Euroscarf |
| BY4741-*trm2Δ* | BY4741, *YKR056w::kanMX* | Euroscarf |
| BY4741-*trm4Δ* | BY4741, *YBL024w::kanMX* | Euroscarf |
| BY4741-*trm8Δ* | BY4741, *YDL201w::kanMX* | Euroscarf |
| BY4741-*trm11Δ* | BY4741, *YOL124c::kanMX* | Euroscarf |
| c13-ABYS-86 | *MATα ura3Δ5 leu2-3,112 his3 pra1-1 prb1-1 prc1-1 cps1-3* (wild-type) | (4) |
| c13-ABYS-86-*dus1Δ* | c13-ABYS-86, *YML080w::kanMX* | This study |
| c13-ABYS-86-*pus4Δ* | c13-ABYS-86, *YNL292w::kanMX* | This study |
| c13-ABYS-86-*trm1Δ* | c13-ABYS-86, *YDR120c::kanMX* | This study |
| c13-ABYS-86-*trm2Δ* | c13-ABYS-86, *YKR056w::kanMX* | This study |
| c13-ABYS-86-*trm4Δ* | c13-ABYS-86, *YBL024w::kanMX* | This study |
| c13-ABYS-86-*trm8Δ* | c13-ABYS-86, *YDL201w::kanMX* | This study |
| c13-ABYS-86-*trm11Δ* | c13-ABYS-86, *YOL124c::kanMX* | This study |

**Supplementary Table S3: Mass spectrometric parameters for detection of RNA nucleosides by isotope dilution mass spectrometry.**

| Compound Group | Compound Name | Precursor Ion (m/z) | Product Ion (m/z) | Ret Time (min) | Delta Ret Time (min) | Fragmentor (V) | Collision Energy (eV) | Cell Accelerator Voltage (V) | Polarity |
|---|---|---|---|---|---|---|---|---|---|
| unlabelled | C | 244.1 | 112.1 | 2.1 | 1 | 175 | 13 | 5 | Positive |
| | U | 245.1 | 113.1 | 2.9 | 1 | 95 | 5 | 5 | Positive |
| | G | 284.1 | 152.0 | 4.1 | 1 | 150 | 17 | 5 | Positive |
| | A | 268.1 | 136.1 | 5.2 | 1 | 200 | 25 | 5 | Positive |
| | D | 247.1 | 115.0 | 1.7 | 1 | 70 | 5 | 5 | Positive |
| | Ψ | 245.1 | 209.0 | 1.7 | 1 | 90 | 5 | 5 | Positive |
| | $m^1A$ | 282.1 | 150.0 | 3.3 | 2 | 150 | 25 | 5 | Positive |
| | $m^5C$ | 258.1 | 126.1 | 3.5 | 1 | 185 | 13 | 5 | Positive |
| | Cm | 258.0 | 112.0 | 3.8 | 1 | 180 | 9 | 5 | Positive |
| | $m^7G$ | 298.1 | 166.0 | 3.8 | 1 | 100 | 13 | 5 | Positive |
| | I | 269.1 | 137.0 | 4.0 | 1 | 100 | 10 | 5 | Positive |
| | $m^5U$ | 259.1 | 127.1 | 4.1 | 1 | 95 | 9 | 5 | Positive |
| | Gm | 298.1 | 152.0 | 4.9 | 1 | 100 | 9 | 5 | Positive |
| | $m^2G$ | 298.1 | 166.0 | 5.0 | 1 | 95 | 17 | 5 | Positive |
| | $m^2_2G$ | 312.1 | 180.0 | 5.6 | 1 | 130 | 13 | 5 | Positive |
| | $m^1G$ | 298.1 | 166.0 | 4.8 | 1 | 130 | 13 | 5 | Positive |
| | $m^6A$ | 282.1 | 150.0 | 6.5 | 1 | 140 | 17 | 5 | Positive |
| SILIS | C SILIS | 253.0 | 116.0 | 2.1 | 1 | 175 | 13 | 5 | Positive |
| | U SILIS | 254.0 | 117.0 | 2.9 | 1 | 95 | 5 | 5 | Positive |
| | G SILIS | 294.0 | 157.0 | 4.1 | 1 | 150 | 17 | 5 | Positive |
| | A SILIS | 278.0 | 141.0 | 5.2 | 1 | 200 | 25 | 5 | Positive |
| | D SILIS | 256.0 | 119.0 | 1.7 | 1 | 70 | 5 | 5 | Positive |
| | Ψ SILIS | 254.0 | 218.0 | 1.7 | 1 | 90 | 5 | 5 | Positive |
| | $m^1A$ SILIS | 295.0 | 158.0 | 3.3 | 2 | 150 | 25 | 5 | Positive |
| | $m^5C$ SILIS | 270.0 | 133.0 | 3.5 | 1 | 185 | 13 | 5 | Positive |
| | Cm SILIS | 270.0 | 116.0 | 3.8 | 1 | 180 | 9 | 5 | Positive |
| | $m^7G$ SILIS | 311.0 | 174.0 | 3.8 | 1 | 100 | 13 | 5 | Positive |
| | I SILIS | 279.0 | 142.0 | 4.0 | 1 | 100 | 10 | 5 | Positive |
| | $m^5U$ SILIS | 271.0 | 134.0 | 4.1 | 1 | 95 | 9 | 5 | Positive |
| | Gm SILIS | 311.0 | 157.0 | 4.9 | 1 | 100 | 9 | 5 | Positive |
| | $m^2G$ SILIS | 311.0 | 174.0 | 5.0 | 1 | 95 | 17 | 5 | Positive |
| | $m^2_2G$ SILIS | 328.0 | 191.0 | 5.6 | 1 | 130 | 13 | 5 | Positive |
| | $m^1G$ SILIS | 311.0 | 174.0 | 4.8 | 1 | 130 | 13 | 5 | Positive |
| | $m^6A$ SILIS | 295.0 | 158.0 | 6.5 | 1 | 140 | 17 | 5 | Positive |

Abbreviations: C, cytidine; U, uridine; G, guanosine; A, adenosine; D, dihydrouridine; Ψ, pseudouridine; $m^1A$, 1-methyladenosine; $m^5C$, 5-methylcytidine; Cm, 2'OMe-cytidine; $m^7G$, 7-methylguanosine; I, inosine; $m^5U$, 5-methyluridine; Gm, 2'OMe-guanosine; $m^2G$, 2-methylguanosine; $m^2_2G$, 2,2-dimethylguanosine; $m^1G$, 1-methylguanosine; and $m^6A$, 6-methyladenosine. SILIS: stable isotope labelled internal standard.

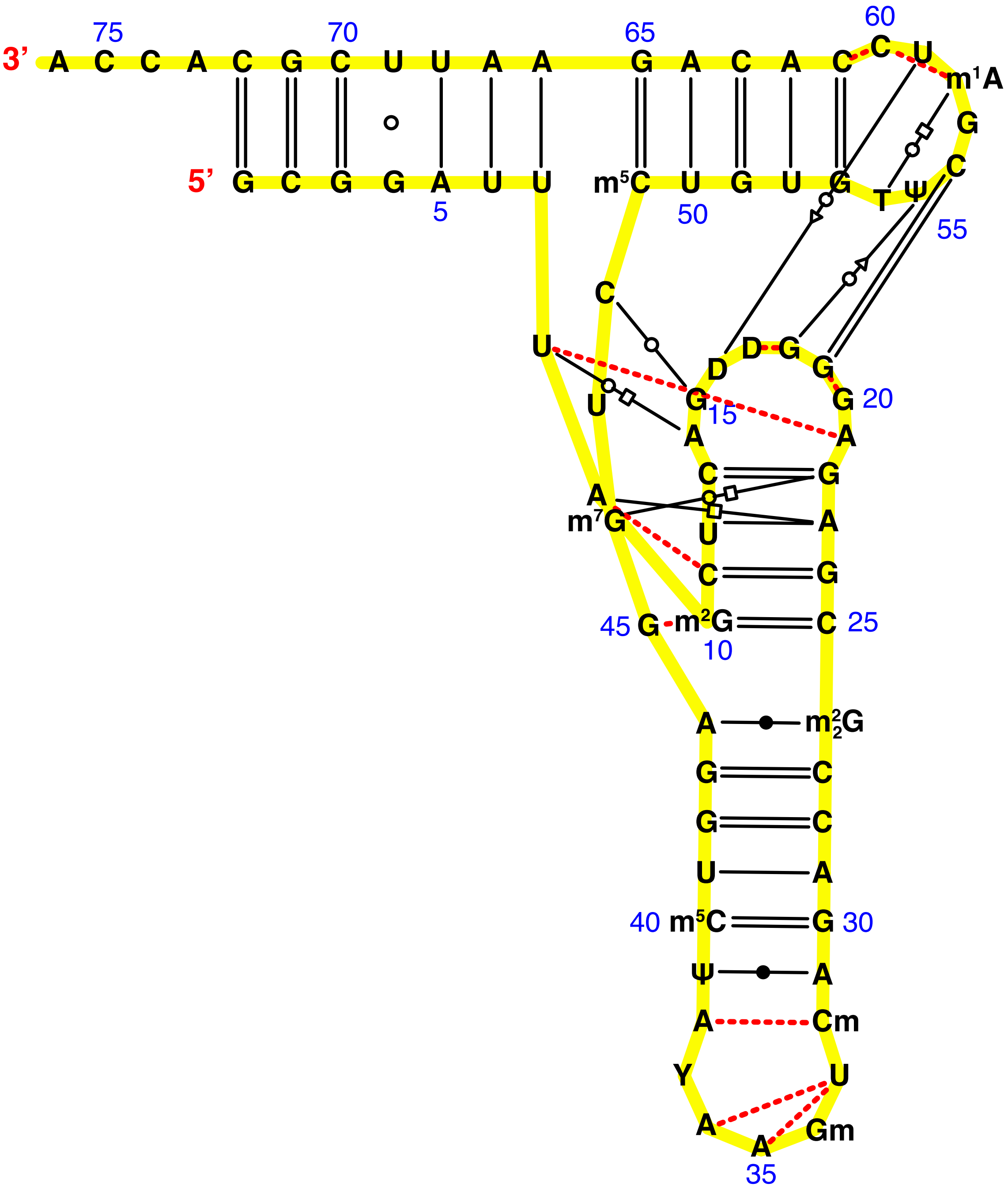


**Supplementary Figure S1: Three-dimensional L-shaped structure of yeast tRNA$^{Phe}$.**

The L-shaped representation is based on the analysis of the PDB coordinates of yeast tRNA$^{Phe}$ (PDB code 1ehz) (5) with the program RNAMLview (6). The standard Watson-Crick base pairs are annotated with two parallel lines for GC pairs and a single line for AU pairs. GU wobble pairs are annotated with a circle. Single dashed lines mean interactions formed *via* a single hydrogen bond. All the other 12 type of non-Watson-Crick base pairs are annotated with symbols according to the classification defined by Leontis and Westhof (7).

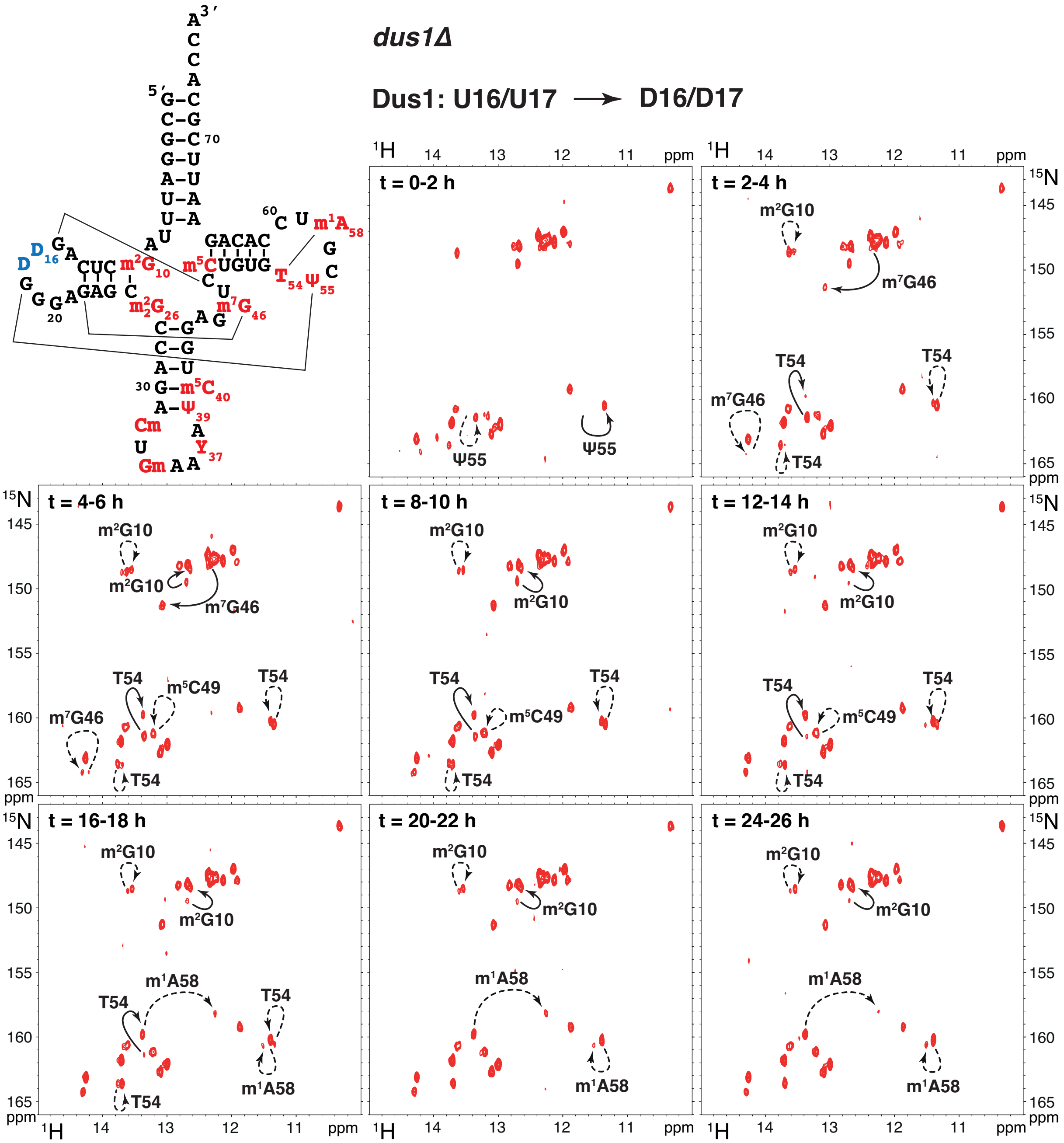


**Supplementary Figure S2: Time-resolved NMR monitoring of yeast tRNA$^{Phe}$ maturation in *dus1Δ* yeast extracts.**

(*top left*) Sequence and cloverleaf representation of modified yeast tRNA$^{Phe}$. Main tertiary interactions are represented with thin lines. Modifications are in red. The modifications introduced by the Dus1 enzyme and missing in these extracts are in blue.

(*spectra from top to bottom right*) Imino ($^{1}$H,$^{15}$N) correlation spectra of a $^{15}$N-labelled tRNA$^{Phe}$ measured in a time-resolved fashion during a continuous incubation at 30°C in yeast extract from a *dus1Δ* strain over 26 h. Each NMR spectrum measurement corresponds to a 2 hour time period, as indicated on the top-left corner of each spectrum. Modifications occurring at the different steps are reported with plain arrows for direct effects, or dashed arrows for indirect effects.

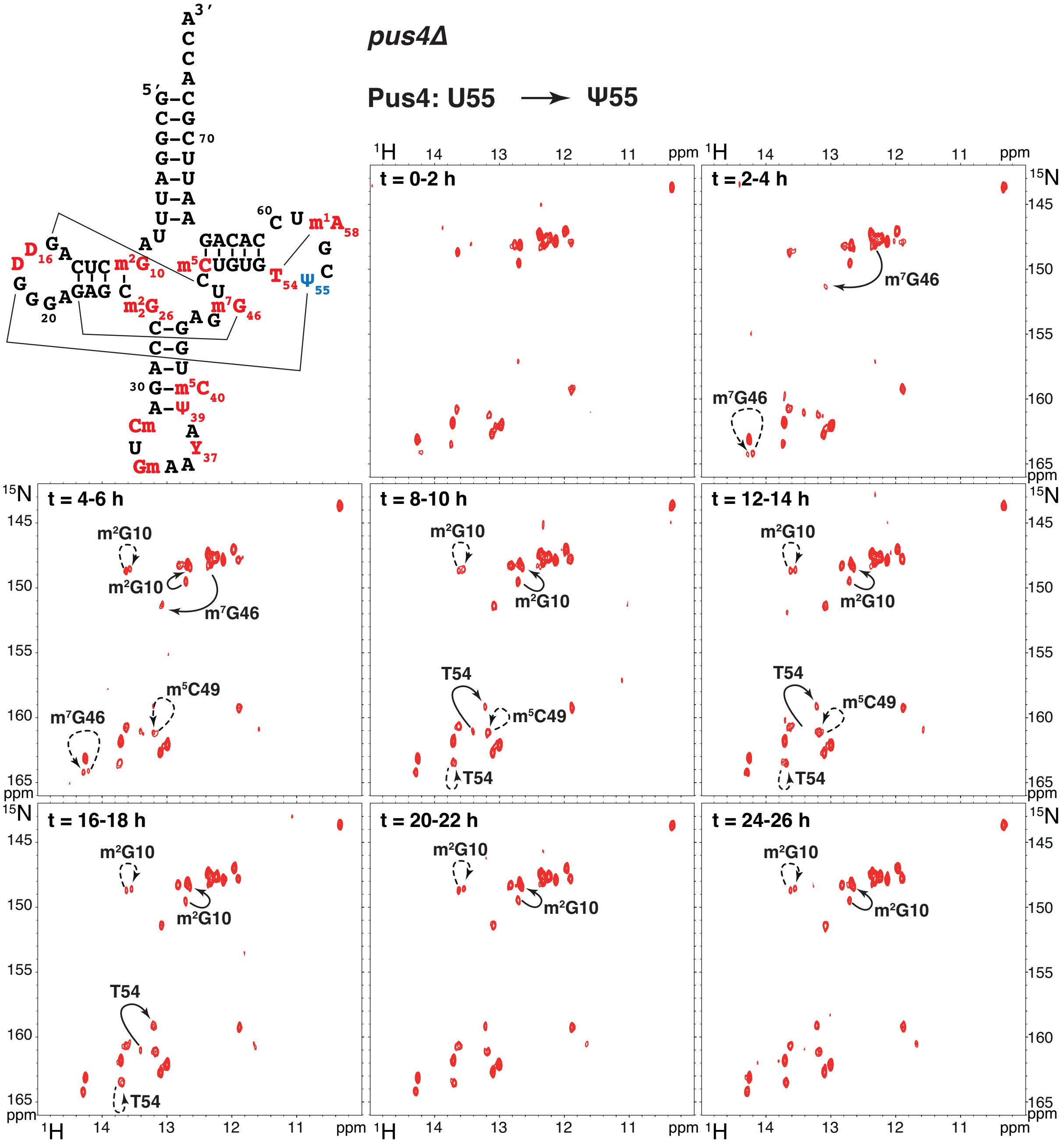


**Supplementary Figure S3: Time-resolved NMR monitoring of yeast tRNA$^{Phe}$ maturation in *pus4Δ* yeast extracts.**

(*top left*) Sequence and cloverleaf representation of modified yeast tRNA$^{Phe}$. Main tertiary interactions are represented with thin lines. Modifications are in red. The modification introduced by the Pus4 enzyme and missing in these extracts is in blue.

(*spectra from top to bottom right*) Imino ($^{1}H$,$^{15}N$) correlation spectra of a $^{15}N$-labelled tRNA$^{Phe}$ measured in a time-resolved fashion during a continuous incubation at 30°C in yeast extract from a *pus4Δ* strain over 26 h. Each NMR spectrum measurement corresponds to a 2 hour time period, as indicated on the top-left corner of each spectrum. Modifications occurring at the different steps are reported with plain arrows for direct effects, or dashed arrows for indirect effects.

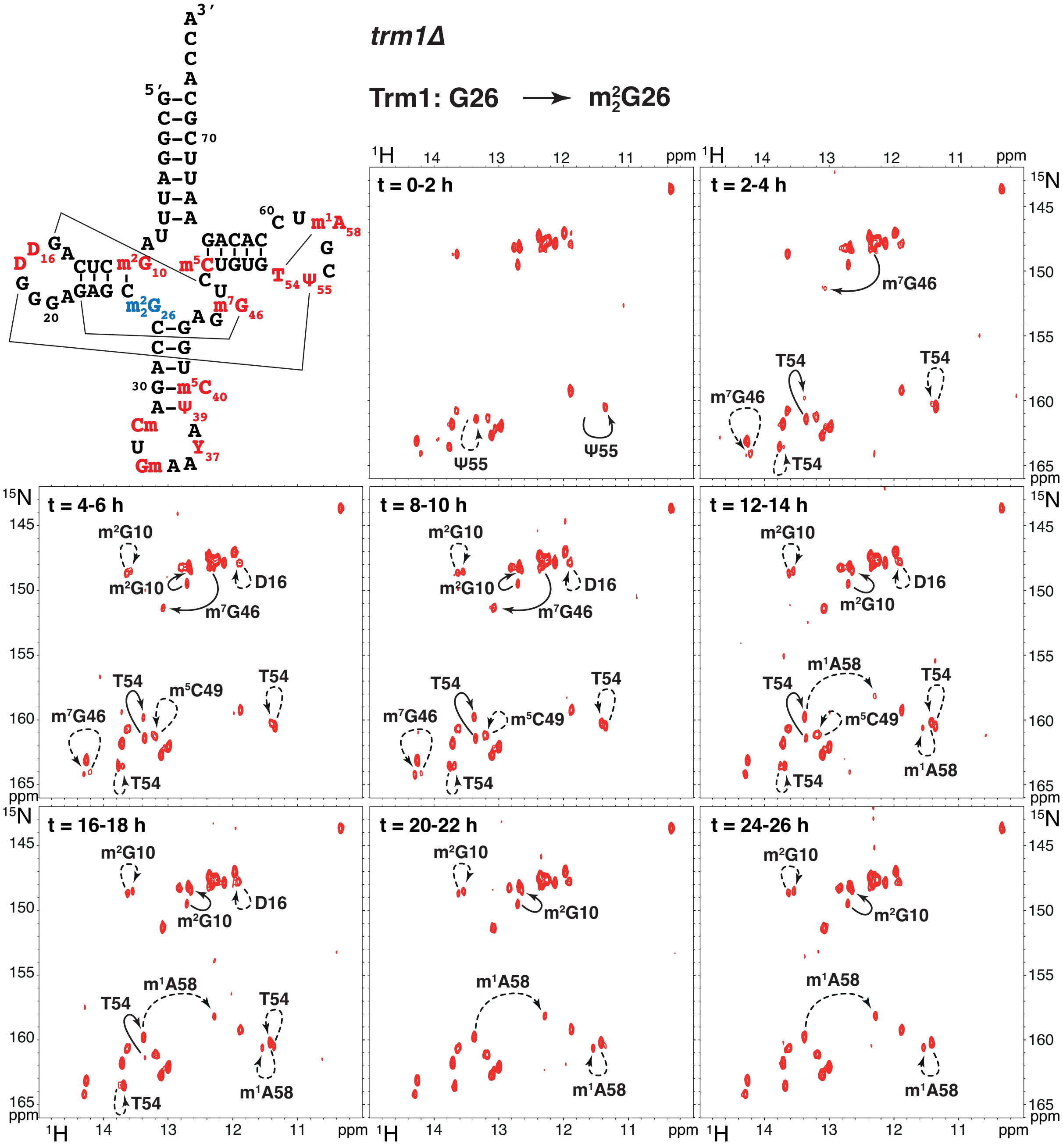


**Supplementary Figure S4: Time-resolved NMR monitoring of yeast $tRNA^{Phe}$ maturation in *trm1Δ* yeast extracts.**

(*top left*) Sequence and cloverleaf representation of modified yeast $tRNA^{Phe}$. Main tertiary interactions are represented with thin lines. Modifications are in red. The modification introduced by the Trm1 enzyme and missing in these extracts is in blue.

(*spectra from top to bottom right*) Imino ($^1H$,$^{15}N$) correlation spectra of a $^{15}N$-labelled $tRNA^{Phe}$ measured in a time-resolved fashion during a continuous incubation at 30°C in yeast extract from a *trm1Δ* strain over 26 h. Each NMR spectrum measurement corresponds to a 2 hour time period, as indicated on the top-left corner of each spectrum. Modifications occurring at the different steps are reported with plain arrows for direct effects, or dashed arrows for indirect effects.

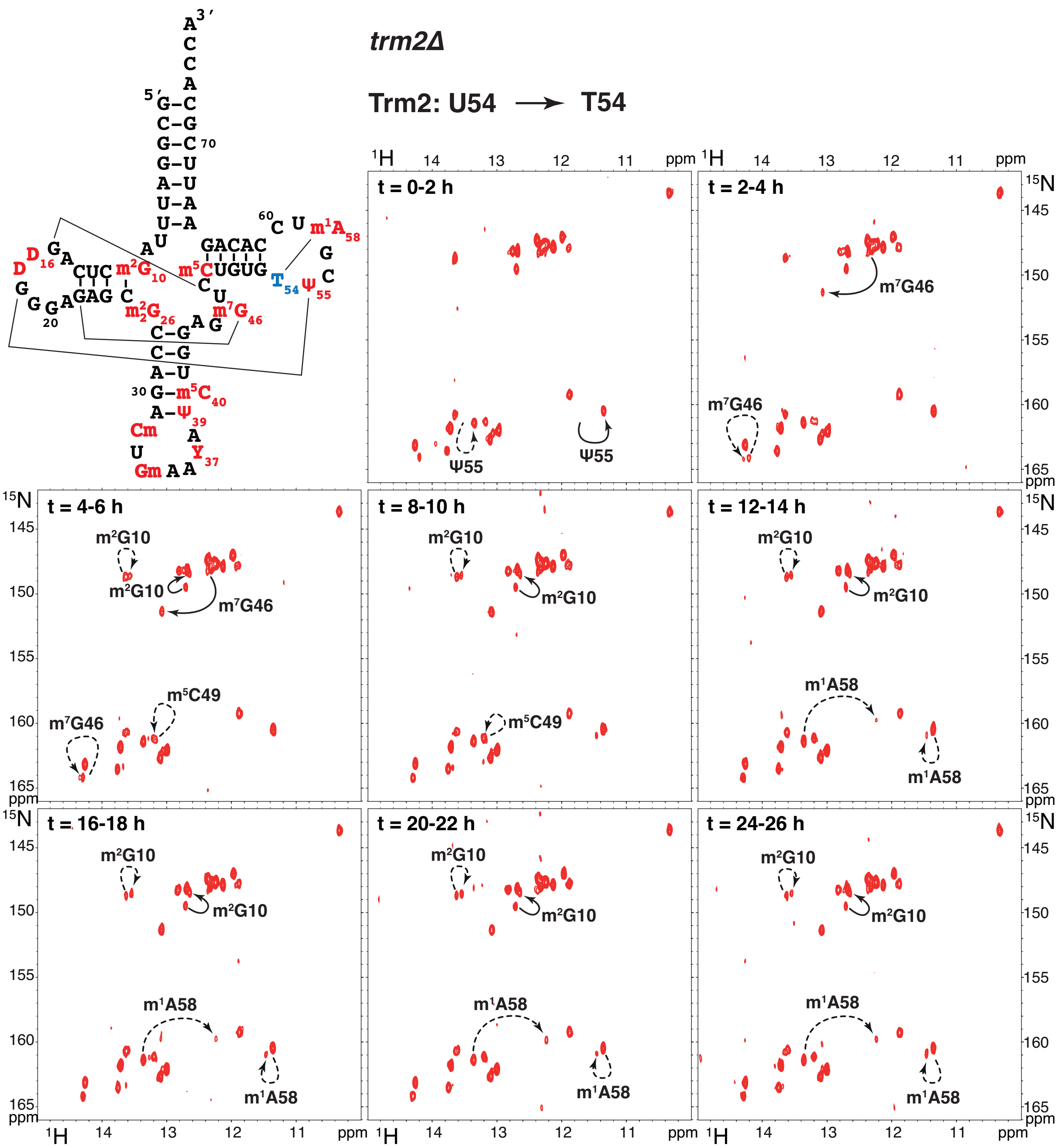


**Supplementary Figure S5: Time-resolved NMR monitoring of yeast tRNA$^{Phe}$ maturation in *trm2Δ* yeast extracts.**

(*top left*) Sequence and cloverleaf representation of modified yeast tRNA$^{Phe}$. Main tertiary interactions are represented with thin lines. Modifications are in red. The modification introduced by the Trm2 enzyme and missing in these extracts is in blue.

(*spectra from top to bottom right*) Imino ($^{1}$H,$^{15}$N) correlation spectra of a $^{15}$N-labelled tRNA$^{Phe}$ measured in a time-resolved fashion during a continuous incubation at 30°C in yeast extract from a *trm2Δ* strain over 26 h. Each NMR spectrum measurement corresponds to a 2 hour time period, as indicated on the top-left corner of each spectrum. Modifications occurring at the different steps are reported with plain arrows for direct effects, or dashed arrows for indirect effects.

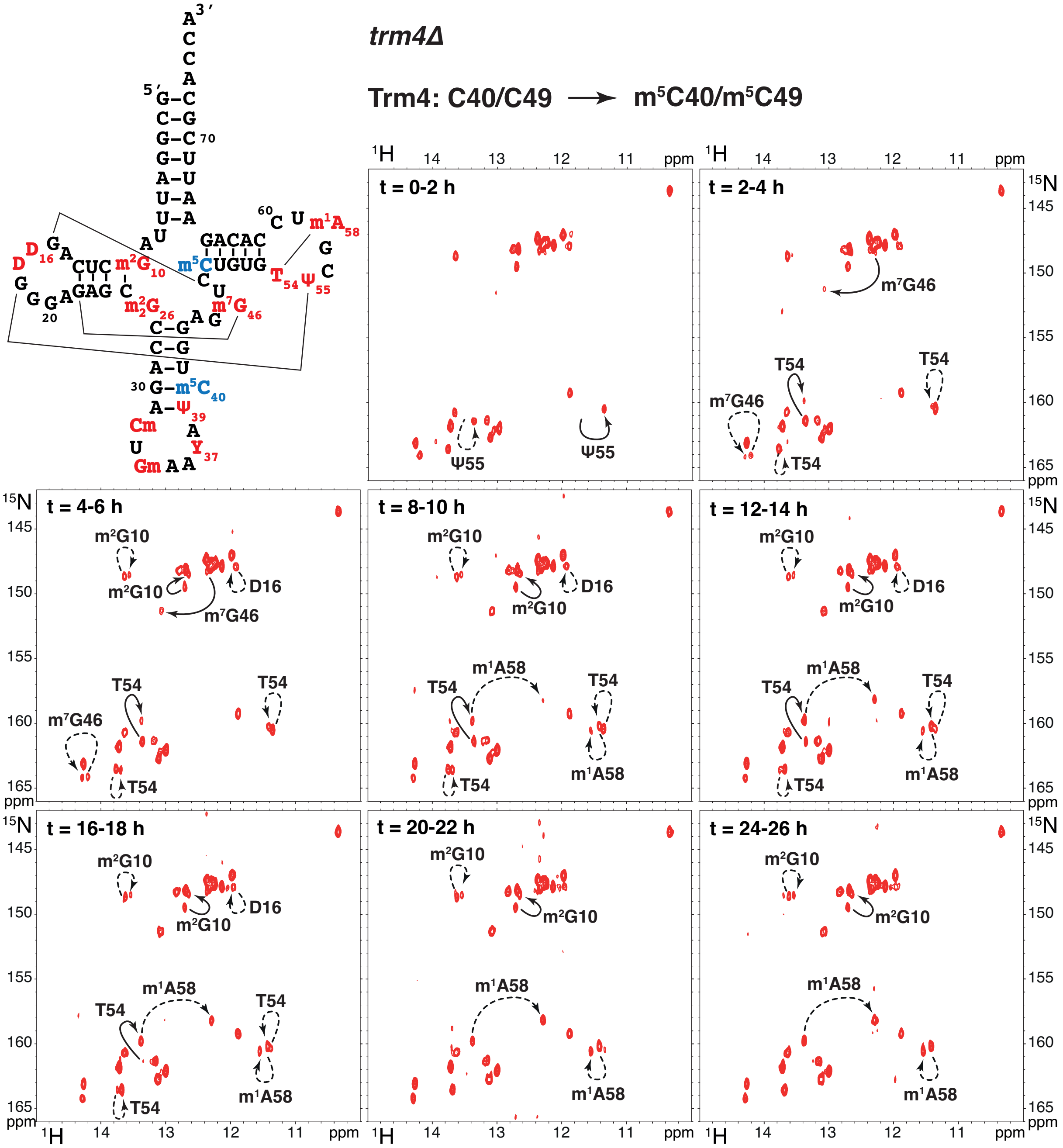


**Supplementary Figure S6: Time-resolved NMR monitoring of yeast tRNA$^{Phe}$ maturation in *trm4Δ* yeast extracts.**

(*top left*) Sequence and cloverleaf representation of modified yeast tRNA$^{Phe}$. Main tertiary interactions are represented with thin lines. Modifications are in red. The modifications introduced by the Trm4 enzyme and missing in these extracts are in blue.

(*spectra from top to bottom right*) Imino ($^{1}$H,$^{15}$N) correlation spectra of a $^{15}$N-labelled tRNA$^{Phe}$ measured in a time-resolved fashion during a continuous incubation at 30°C in yeast extract from a *trm4Δ* strain over 26 h. Each NMR spectrum measurement corresponds to a 2 hour time period, as indicated on the top-left corner of each spectrum. Modifications occurring at the different steps are reported with plain arrows for direct effects, or dashed arrows for indirect effects.

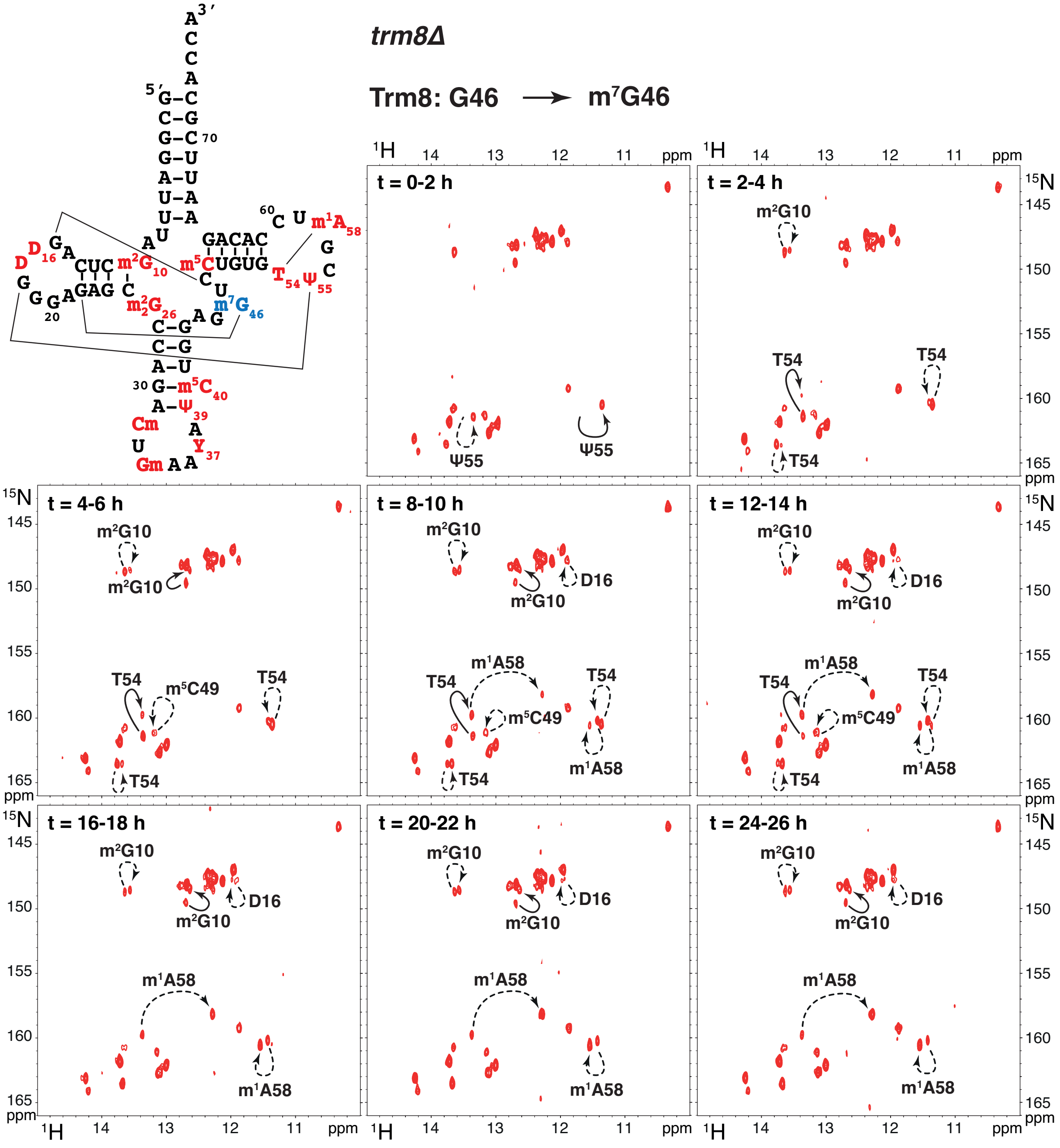


**Supplementary Figure S7: Time-resolved NMR monitoring of yeast tRNA$^{Phe}$ maturation in *trm8Δ* yeast extracts.**

(*top left*) Sequence and cloverleaf representation of modified yeast tRNA$^{Phe}$. Main tertiary interactions are represented with thin lines. Modifications are in red. The modification introduced by the Trm8 enzyme and missing in these extracts is in blue.

(*spectra from top to bottom right*) Imino ($^{1}$H,$^{15}$N) correlation spectra of a $^{15}$N-labelled tRNA$^{Phe}$ measured in a time-resolved fashion during a continuous incubation at 30°C in yeast extract from a *trm8Δ* strain over 26 h. Each NMR spectrum measurement corresponds to a 2 hour time period, as indicated on the top-left corner of each spectrum. Modifications occurring at the different steps are reported with plain arrows for direct effects, or dashed arrows for indirect effects.

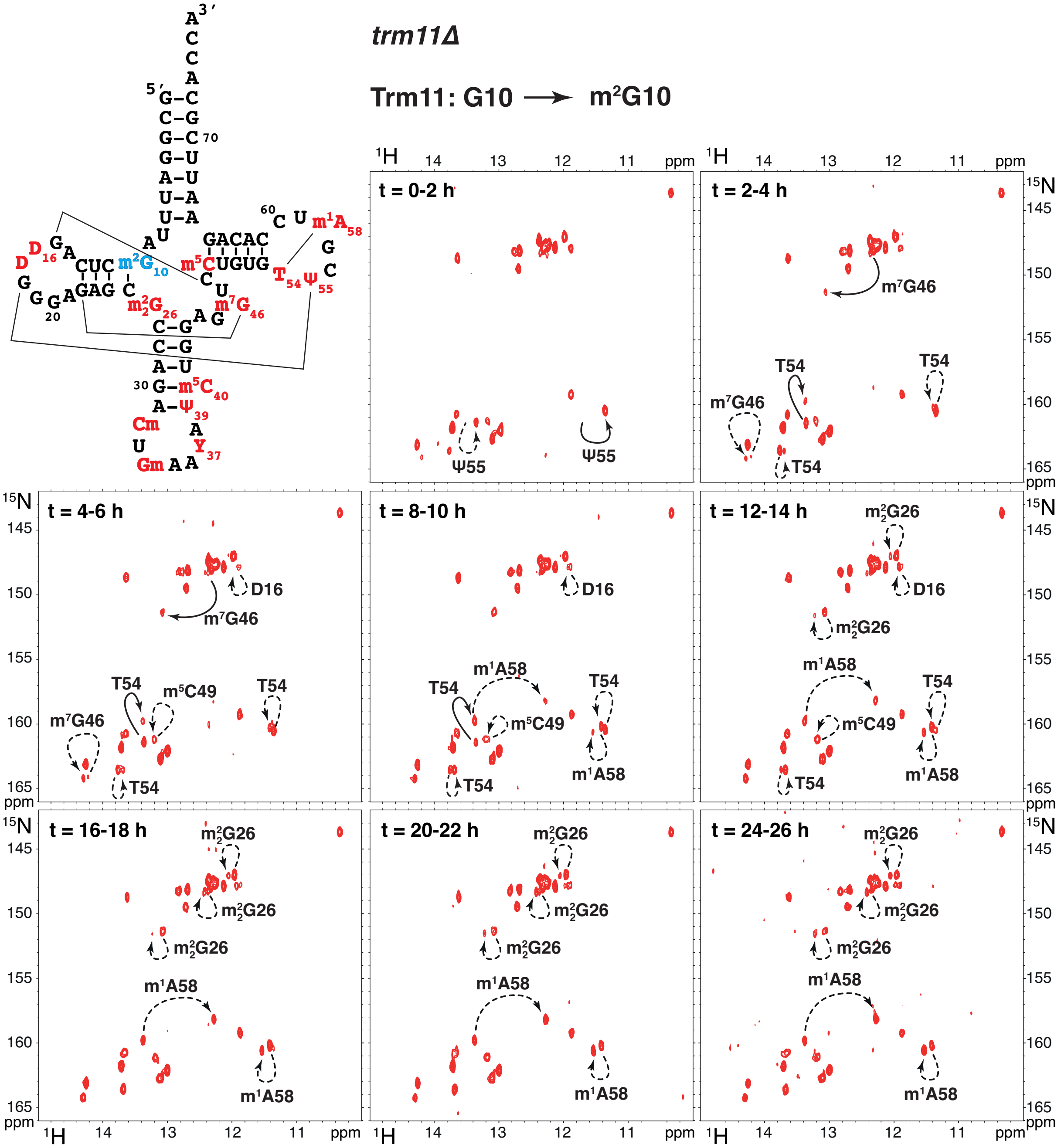


**Supplementary Figure S8: Time-resolved NMR monitoring of yeast tRNA$^{Phe}$ maturation in *trm11Δ* yeast extracts.**

(*top left*) Sequence and cloverleaf representation of modified yeast tRNA$^{Phe}$. Main tertiary interactions are represented with thin lines. Modifications are in red. The modification introduced by the Trm11 enzyme and missing in these extracts is in blue.

(*spectra from top to bottom right*) Imino ($^1$H,$^{15}$N) correlation spectra of a $^{15}$N-labelled tRNA$^{Phe}$ measured in a time-resolved fashion during a continuous incubation at 30°C in yeast extract from a *trm11Δ* strain over 26 h. Each NMR spectrum measurement corresponds to a 2 hour time period, as indicated on the top-left corner of each spectrum. Modifications occurring at the different steps are reported with plain arrows for direct effects, or dashed arrows for indirect effects.

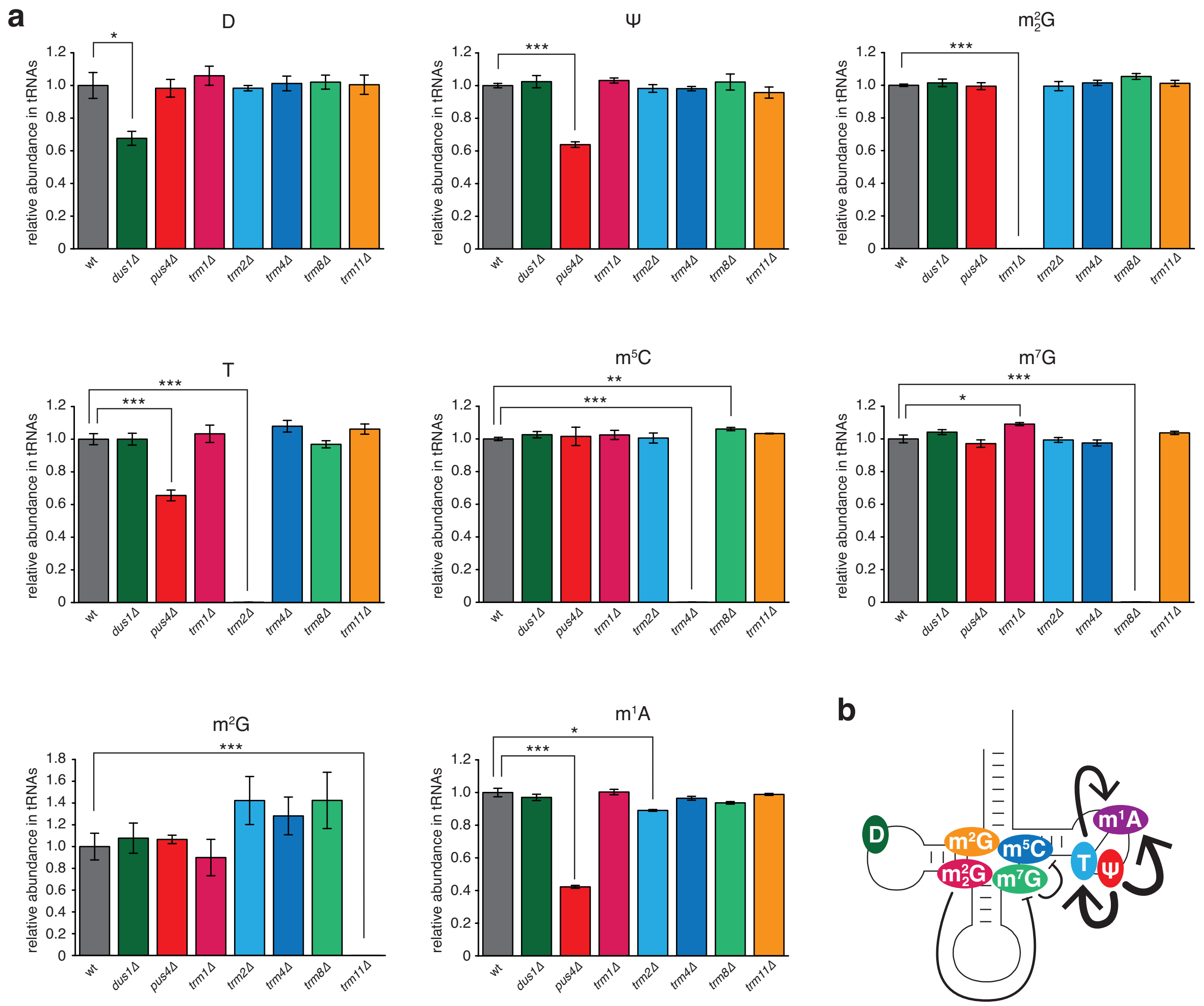


**Supplementary Figure S9: Quantitative analysis of nucleoside modifications in yeast total tRNAs with LC-MS/MS.**

(**a**) Histograms showing the relative abundance of D, Ψ, $m^2_2G$, T, $m^5C$, $m^7G$, $m^2G$, and $m^1A$ in total yeast tRNAs prepared from depleted strains (*dus1Δ*, *pus4Δ*, *trm1Δ*, *trm2Δ*, *trm4Δ*, *trm8Δ* and *trm11Δ*) using the wild-type levels as reference. Significant changes compared to wild type are reported as *** for $p<0.001$, ** for $p< 0.01$ and * for $p<0.05$, n=3. All other changes are not statistically significant. (**b**) Schematic view of the modification circuits revealed by the MS quantification of modifications in yeast total tRNAs. Each modification is displayed on the cloverleaf structure with its associated color. Arrows indicate stimulatory effects and blunted lines inhibitory effects. Thick and thin lines indicate strong and weak effects, respectively.

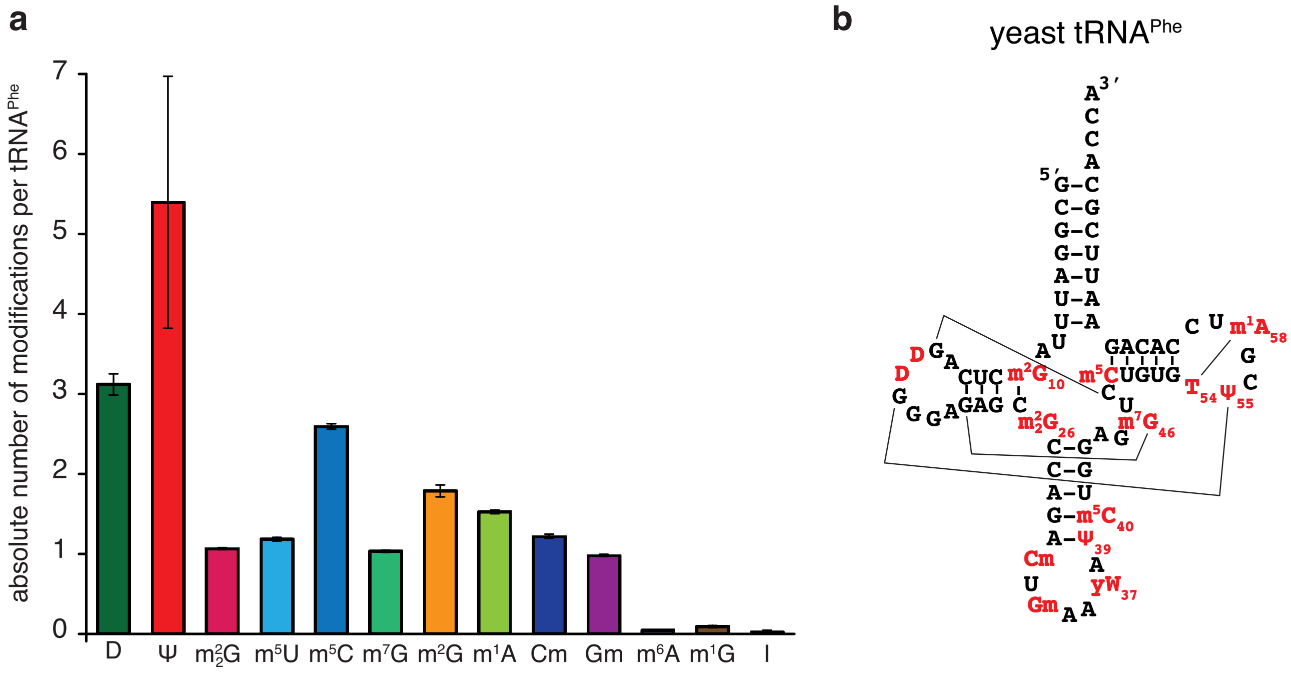


**Supplementary Figure S10: Absolute quantification of nucleoside modifications in specifically purified yeast tRNA$^{Phe}$ with LC-MS/MS.**

(**a**) Histograms showing the absolute abundance of modified nucleosides in tRNA$^{Phe}$ isolated from wild-type yeast. Due to chromatographic separation issues and low mass spectrometric detection efficiency, the quantification of pseudouridine is disturbed, which leads to an overestimation of pseudouridine in the measurements (also reflected in large error bars) (**b**) Sequence and cloverleaf representation of yeast tRNA$^{Phe}$. Modifications are in red.

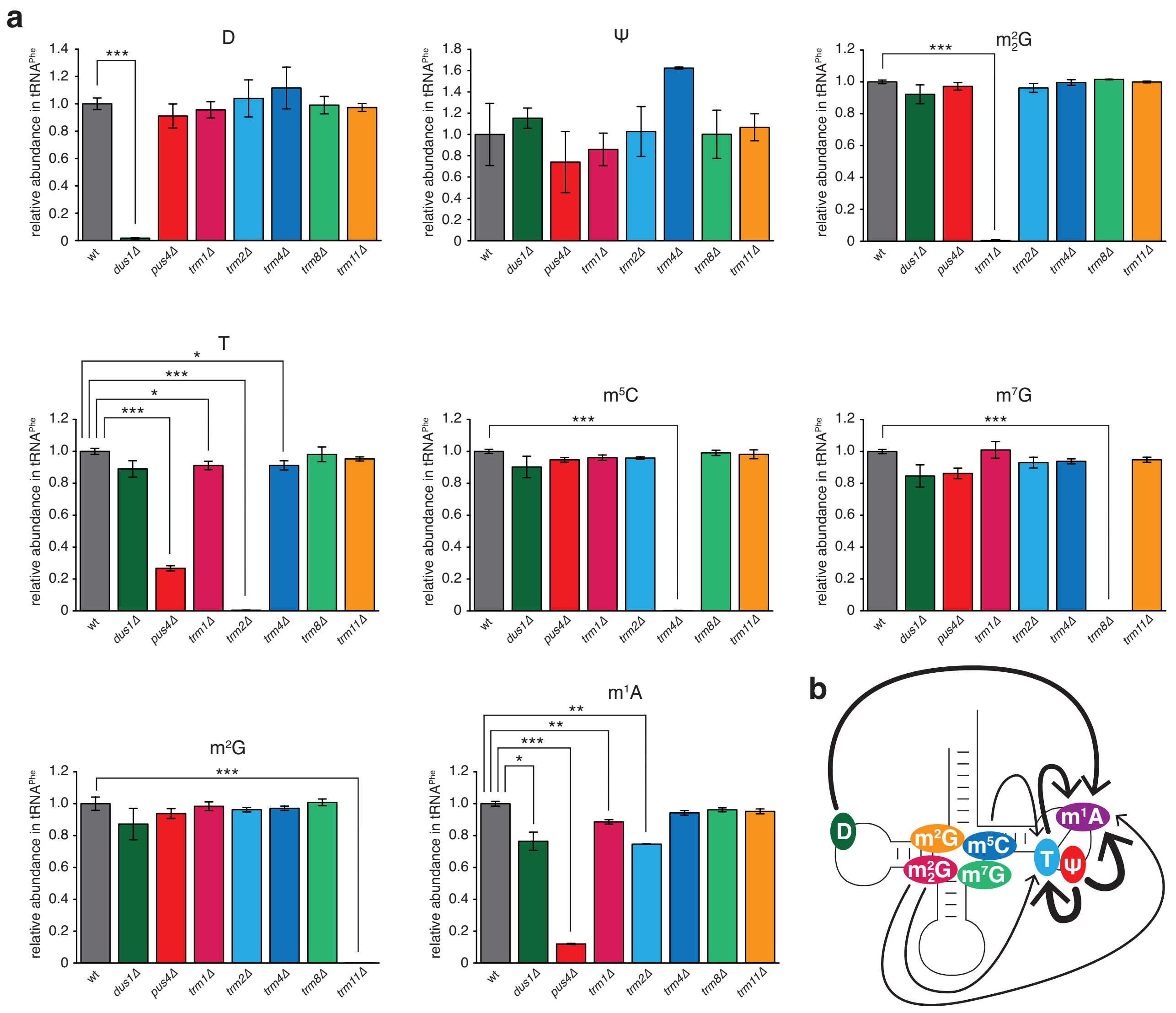


**Supplementary Figure S11: Quantitative analysis of nucleoside modifications in specifically purified yeast tRNA$^{Phe}$ with LC-MS/MS.**

(**a**) Histograms showing the relative abundance of D, Ψ, $m^2_2G$, T, $m^5C$, $m^7G$, $m^2G$, and $m^1A$ in specifically purified yeast tRNA$^{Phe}$ prepared from depleted strains (*dus1Δ*, *pus4Δ*, *trm1Δ*, *trm2Δ*, *trm4Δ*, *trm8Δ* and *trm11Δ*) using the wild-type levels as reference. Significant changes compared to wild type are reported as *** for p<0.001, ** for p< 0.01 and * for p<0.05, n=3. All other changes are not statistically significant. (**b**) Schematic view of the modification circuits revealed by the MS quantification of modifications in specifically purified yeast tRNA$^{Phe}$. Each modification is displayed on the cloverleaf structure with its associated color. Arrows indicate stimulatory effects. Thick and thin lines indicate strong and weak effects, respectively.
